# Programming effects of peripubertal stress on spatial learning

**DOI:** 10.1101/2020.09.22.307702

**Authors:** S. Tzanoulinou, E. Gantelet, C. Sandi, C. Marquez

**Author notes:** Department of Biomedical Sciences, University of Lausanne, Lausanne, Switzerland (S. Tzanoulinou).

## Abstract

Exposure to adversity during early life can have profound influences on brain function and behavior later in life. The peripubertal period is emerging as an important time-window of susceptibility to stress, with substantial evidence documenting long-term consequences in the emotional and social domains. However, little is known about how stress during this period impacts subsequent cognitive functioning. Here, we assessed potential long-term effects of peripubertal stress on spatial learning and memory using the water maze task. In addition, we interrogated whether individual differences in stress-induced behavioral and endocrine changes are related to the degree of adaptation of the corticosterone response to repeated stressor exposure during the peripubertal period. We found that, when tested at adulthood, peripubertally stressed animals displayed a slower learning rate. Strikingly, the level of spatial orientation in the water maze completed on the last training day was predicted by the degree of adaptation of the recovery -and not the peak-of the corticosterone response to stressor exposure (i.e., plasma levels at 60 min post-stressor) across the peripubertal stress period. In addition, peripubertal stress led to changes in emotional and glucocorticoid reactivity to novelty exposure, as well as in the expression levels of the plasticity molecule PSA-NCAM in the hippocampus. Importantly, by assessing the same endpoints in another peripubertally stressed cohort tested during adolescence, we show that the observed effects at adulthood are the result of a delayed programming manifested at adulthood and not protracted effects of stress. Altogether, our results support the view that the degree of stress-induced adaptation of the hypothalamus-pituitary-adrenal axis responsiveness at the important transitional period of puberty relates to the long-term programming of cognition, behavior and endocrine reactivity.

## Introduction

Exposure to adversity during early life can have profound influences on brain function, behavior and cognition at adulthood (Albrecht et al., 2017; Bolton et al., 2017; Sterlemann et al., 2010; Suri et al., 2013), and the precise developmental timing when stress occurs seems to be critical in determining the precise consequences (Gee and Casey, 2015; Lupien et al., 2009). In addition to the recognized impact of neonatal (Bonapersona et al., 2019; Heim and Nemeroff, 2001; Molet et al., 2014; Veenema, 2009) and childhood/juvenile (Albrecht et al., 2017) stress, the peripubertal period is emerging as a time-window of high vulnerability to the programming of emotional (Cordero et al., 2012; Latsko et al., 2016; Márquez et al., 2013; Sheth et al., 2017) and social (Márquez et al., 2013; Poirier et al., 2014; Tzanoulinou et al., 2014a; 2014b) effects of stress (for a review, see (Tzanoulinou and Sandi, 2017). However, despite the well-known modulatory power of stress on cognition (Lupien et al., 2009; Sandi, 2013), little is known about the impact of peripubertal stress on later life cognitive functioning. A few studies in which stressors were applied during the period expanding from peripuberty till young adulthood have reported enduring learning and memory impairments specifically for the spatial domain (Isgor et al., 2004; Sterlemann et al., 2010). Therefore, whether the peripubertal period *per se* is susceptible to long-term programming effects of stress on spatial learning, while plausible, remains unclear.

The peripubertal period, involving time-windows right before and after puberty, comprises drastic hormonal, neurobiological and behavioral changes (Andersen and Teicher, 2008; Blakemore, 2008; Casey et al., 2010; Paus et al., 2008; Romeo et al., 2016; Spear, 2000; Tzanoulinou and Sandi, 2016). In particular, this period involves marked changes in the responsivity of the hypothalamic-pituitary-adrenal (HPA) axis to stressful experiences (McCormick et al., 2017; Romeo et al., 2016), and this transition can be modified by experiences (Gunnar et al., 2019), particularly stressful ones (Kumsta et al., 2017; Márquez et al., 2013; McCormick et al., 2017; Romeo et al., 2016). Strikingly, individual differences in the adaptation of the glucocorticoid response to repeated stress exposure during the peripubertal period in rats were found to predict subsequent changes in emotional and social phenotypes observed during adolescence (Papilloud et al., 2019) and adulthood (Walker et al., 2018; 2017). In addition, genetic selection in rats for the degree of corticosterone adaptation during peripubertal stress (Walker and Sandi, 2018) underscored genetic line-related differences in spatial learning and memory performance (Huzard et al., 2020). Accordingly, given the strong modulatory capacity of glucocorticoids on brain function and cognition (de Quervain et al., 2017; Sandi, 2011), including spatial learning (Akirav et al., 2004; Conboy et al., 2010; Sandi et al., 1997), we hypothesize that long-term programming of peripubertal stress on spatial learning would depend on the individual degree of glucocorticoid adaptation to repeated stress.

When considering the glucocorticoid adaptation to repeated stress, it is important to distinguish between the peak and the recovery phases, as they serve different adaptive functions (Romeo et al., 2016). While peak glucocorticoid levels facilitate physiological processes to deal with immediate challenges (de Kloet et al., 2008; Myers et al., 2014), the recovery phase (i.e., returning to baseline) is key to protect the organism from maladaptive overactivation and to prepare it for eventual new challenges (Karatsoreos and McEwen, 2011). Importantly, the peripubertal period has been reported to set a change in HPA responsivity in both humans and rats, including changes not only in the peak but also in the recovery phases (McCormick et al., 2017). We have previously reported a strong link between the magnitude of adaptation of the peak corticosterone response to repeated stressors given during the peripubertal period in rats and subsequent changes in emotional and social behaviors (Papilloud et al., 2018; Walker et al., 2018; 2017). However, in the context of the current study on spatial learning, we hypothesize that it will be the adaptation of the recovery phase of corticosterone responsiveness that predicts spatial learning. This hypothesis is based on several premises. First, on the crucial roles of the hippocampus in both, spatial learning (Bird and Burgess, 2008) and in providing negative feedback to the HPA axis (Herman and Mueller, 2006; Jacobson and Sapolsky, 1991; Kovács and Makara, 1988) and, thus, impacting on the corticosterone recovery phase. Second, on the high density of corticosteroid receptors present in the hippocampus (de Kloet Frontiers in neuroendocrinology Print1991, n.d.) and their involvement in the HPA axis negative feedback (Reul et al., 1990). Finally, high glucocorticoid levels are known to promote plastic changes in hippocampal structure and function (de Kloet et al., 2018; McEwen et al., 2016), including changes in the expression levels of key plasticity molecules, such as PSA-NCAM (Montaron et al., 2003; Nacher et al., 2004). Importantly, PSA-NCAM-a key post-translational modification of the neural cell adhesion molecule (NCAM)-is critically involved in hippocampal plasticity (Kiss and Muller, 2001) and spatial memory (Bisaz et al., 2009) and modulated by stress (Sandi, 2004).

Therefore, we set this study in rats to assess potential long-term effects of peripubertal stress in spatial learning and memory in the water maze at adulthood, and to investigate whether individual differences in stress-induced changes are related to the adaptation of the corticosterone response (peak vs recovery) to repeated stressor exposure during the peripubertal period. In order to have broader information on the behavioral phenotype for data interpretation, we tested animals in emotional reactivity tasks as well. We also measured plasma corticosterone responses to novelty stress shortly before water maze training to assess both how this response relates to peripubertal corticosterone adaptation and whether it is associated with water maze performance. To understand whether any observed effects at adulthood are the result of a delayed programming or already present shortly after peripubertal stress exposure, we performed a second experiment in which animals were tested during late adolescence. Finally, we assessed levels of the learning and plasticity-related molecule PSA-NCAM in the dentate gyrus (DG) of the hippocampus and, in addition, in the medial amygdala (MeA) as a control region. We found that peripubertally stressed animals had a slower learning rate when tested as adult, but no differences at adolescence. Importantly, those animals that showed less adaptation to stress, as assessed by the corticosterone recovery response during peripubertal stressor exposure, were the ones being more impaired in spatial learning as adults. Interestingly, basal levels of PSA-NCAM expression in the DG were also altered in peripubertally stressed animals, when assessed at adulthood, with those rats showing a suboptimal recovery to stress early in life being the ones that show higher levels of PSA-NCAM in the DG proposing a neurobiological mechanism by which peripubertal stress might be altering the normal maturation of plasticity processes in specific brain regions leading to impaired cognitive performance later in life.

## Materials and Methods

### Animals

Experimental subjects were the male offspring of Wistar Han rats purchased from Charles River Laboratories, France, and bred in our animal facility (n=70). All animals were kept in constant conditions of humidity and temperature (22 ± 1°C) with a 12-h light-dark cycle (lights on at 7:00 AM). Food and water were available *ad libitum*. All the procedures described were conducted in conformity with Swiss National Institutional Guidelines on Animal Experimentation, and approved by a license issued from the Swiss Cantonal Veterinary Office Committee for Animal Experimentation.

### Experimental Design

At weaning (P21), male rats from different litters were distributed into different home cages in groups of two non-siblings, and each cage was randomly assigned to control (CTRL, n=34) or peripubertal stress (STRESS, n=36) conditions. Animals from the STRESS group underwent the peripubertal stress protocol (PPS) starting at P28 (Márquez et al., 2013), and CTRL animals were briefly handled and returned to their home cage. Behavior and hormonal characterizations later in life of the experimental groups were performed at adolescence (P55+) and adulthood (P80+) in independent groups of animals (Figure 1) (Adolescence CTRL n=18; Adolescence STRESS n=18; Adulthood CTRL n=16; Adulthood STRESS n=18). Before behavioral testing, animals were handled for 3 consecutive days to acclimatize to the experimenter and general conditions. Animals were tested in an Open Filed and Novel Object test and their stress response was assessed after a novelty challenge by measuring corticosterone plasmatic levels (see below). Then, animals of each experimental group were further divided into two groups, one which would undergo behavioral evaluation of learning and memory in the Morris Water maze (Adolescence CTRL n=10; Adolescence STRESS n=10; Adulthood CTRL n=8; Adulthood STRESS n=10) and a second one that would be used to study basal levels of polysialylated-neural cell adhesion molecule (PSA-NCAM) in specific brain regions by immunohistochemistry in either adolescence or adulthood (Adolescence CTRL n=8; Adolescence STRESS n=8; Adulthood CTRL n=8; Adulthood STRESS n=8; with one adult animal being excluded from one PSA-NCAM measurement due to poor IHC signal due to quality of the tissue). Animals were sacrificed in basal conditions by transcardial perfusion under anesthesia, brains rapidly removed, post fixed in PFA 4% for 4 hours and maintained in PBS until further processing for PSA-NCAM immunohistochemistry.

**Figure 1.**
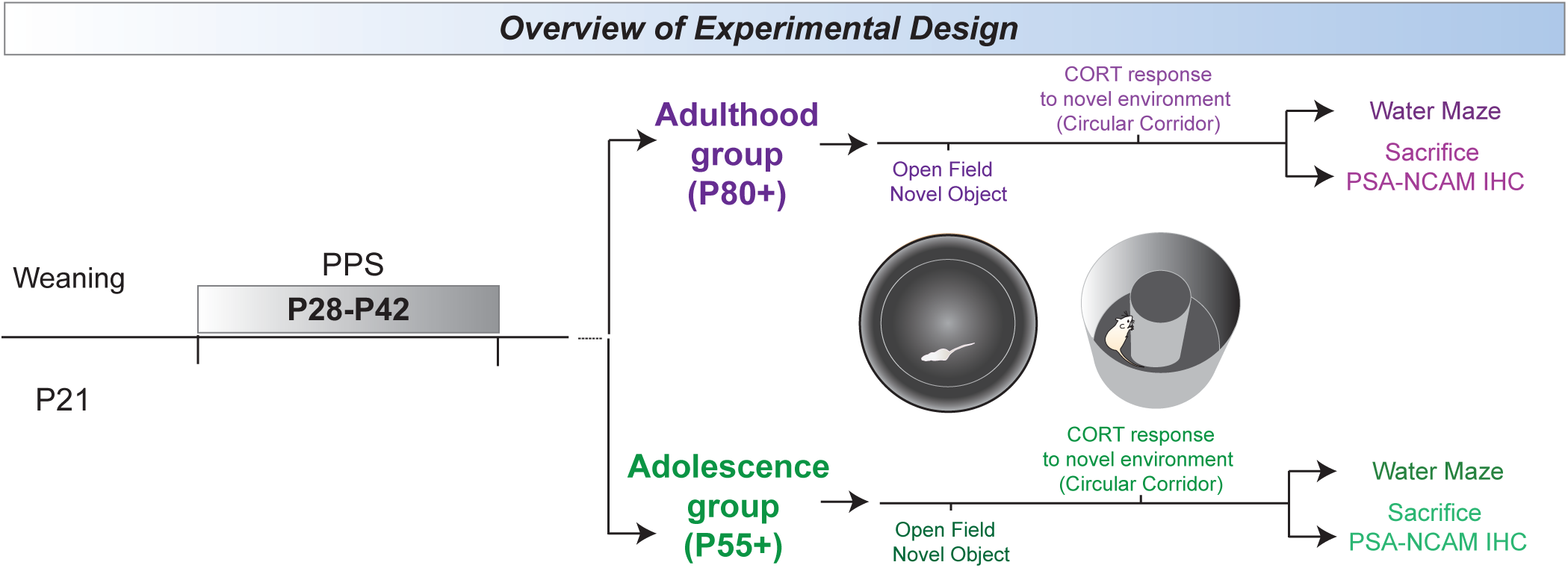
Overview of experimental design to assess the long-term effects of peripubertal stress (PPS) in different moments of development. Rats were weaned at P21 and were either exposed to the PPS protocol from P28 to P42, or assigned to the Control group. The stressors used were exposure to an elevated platform and to a predator odor (TMT) (for more details, please see materials and methods). Control rats were briefly handled on the days of the PPS and then returned to their home cages. Subsequently, control (CTRL) and peripubertally stressed (STRESS) rats were split in two age groups: the adulthood group, and as a control, the adolescence group, depending on when they underwent further tests. All animals were subjected to an open field and novel object exploration tests. Subsequently, their corticosterone reactivity was evaluated after exposure to a novel environment (i.e. exposure to a circular corridor). They were then further split into a group that performed the water maze and a group that was assessed for PSA-NCAM expression levels in the dentate gyrus and medial amygdala.

### Peripubertal Stress Protocol

Peripubertal Stress Protocol (PPS) was performed as previously described (Márquez et al., 2013; Veenit et al., 2014). Specifically, the stress protocol consisted of presenting two different fear-inducing stressors (each one lasting 25 min): (1) exposure to the synthetic fox odor trimethylthiazoline (9 μl) (Phero Tech Inc., Delta, BC, Canada) released through a small cloth, in a plastic box (38 cm length, 27.5 cm width and 31 cm height) placed under a bright light (210–250 lx); and (2) exposure to an elevated platform (12 × 12 cm, elevated 95 cm from the ground) under direct bright light (470–500 lx). The stressors were applied subchronically during the peripubertal period (a total of 7 days across postnatal day P28 to P42, i.e., on P28– P30, P34, P36, P40 and P42), during the light phase, and according to a variable schedule, where the order and timing of the stressors were changed on different days (Figure 2A). On some stress days, only one stressor was presented, while on other days, the two stressors were given consecutively. Following each stress session, animals were returned to their home-cages where a transparent Plexiglas wall with holes separated each animal for 15 minutes before rejoining their cage mates. On the first and last day of Peripubertal stress, blood samples were collected at different time points only to STRESS animals, in order to study the adaptation dynamics to the stress protocol (see below). The control animals were handled on the days that their experimental counterparts were exposed to stress. Animals in the same cage were always assigned to the same experimental group (either CTRL or STRESS).

**Figure 2.**
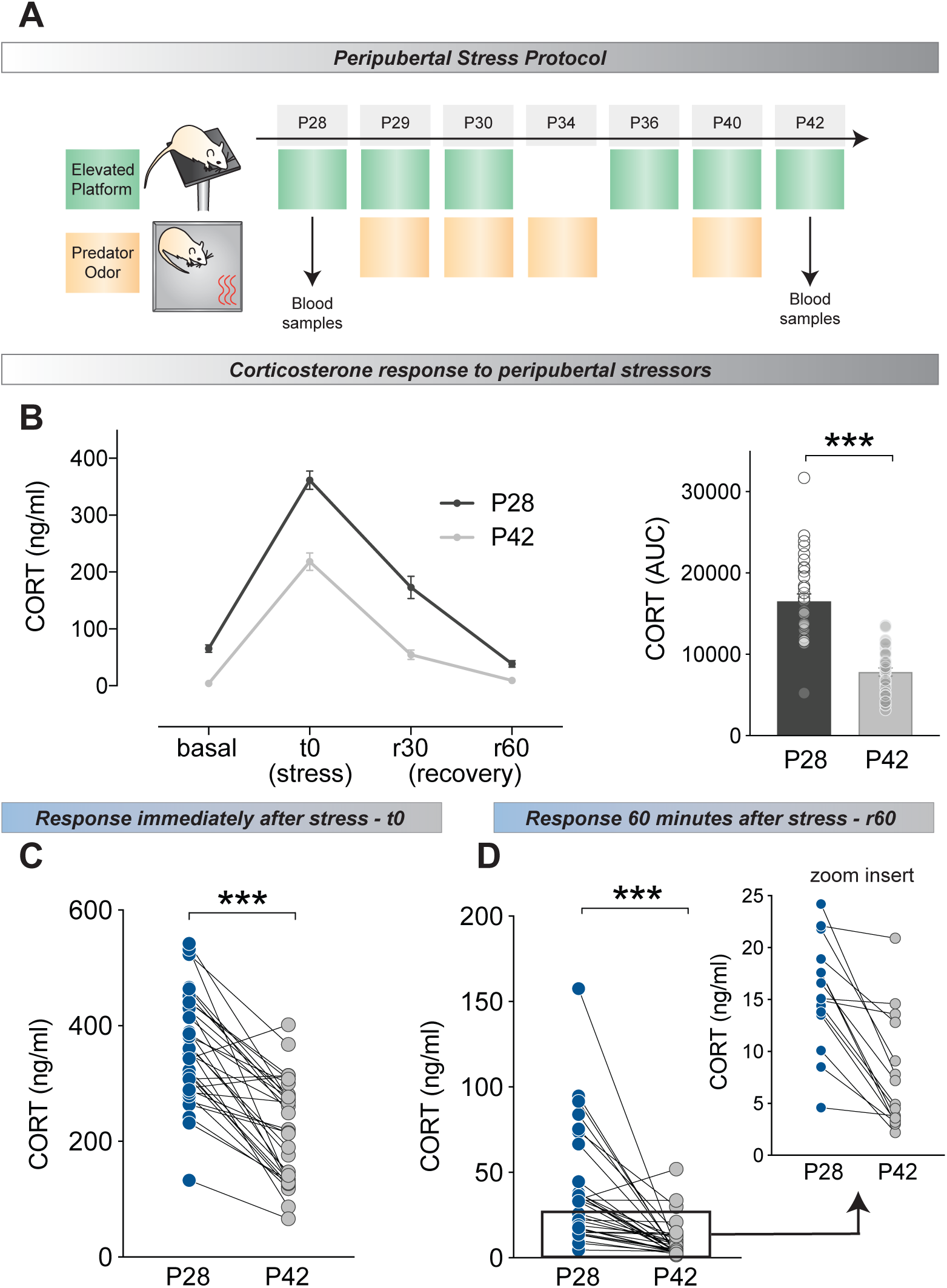
Corticosterone (CORT) response during PPS. **A**. Overview of PPS protocol. Blood samples were obtained on P28 and P42 from STRESS rats at different time points to assess HPA axis reactivity and adaptation to the stressors. **B**. Left: samples were taken at baseline conditions (i.e. before stressor exposure-basal), immediately following the stressor (stress, t0), as well as, 30 minutes and 60 minutes after the end of the exposure to stress (recovery (r) 30 and r60 respectively). Reduced CORT response was observed on P42 as compared to P28. Right: This reduced response is reflected in the CORT area under the curve (AUC) levels that was calculated by considering all four time-points (basal, stress, rec 30 and rec 60). P42 CORT levels correspond to smaller AUC overall. **C**. Individual differences in the ability to adapt to peak (t0) CORT response in P28 and P42 is shown for all animals. Overall, responses to the elevated platform where reduced in P42 compared to the first exposure in P28, however, this decrease was variable among animals. **D**. Variability of adaptation in the recovery period after stress comparing P28 and P42 is shown for all animals. Although CORT levels were low at this time point (60 min after the end of stress exposure) marked individual differences were still observed in the ability to adapt to the subchronic stress. Results are expressed as the mean ± S.E.M. *** p<0.001.

### Open Field and Novel Object Reactivity Tests

Rats’ exploration levels were assessed in the open field test as previously described (Salehi et al., 2010). They were individually placed in the center of the open field arena (a circular open arena with a diameter of 100 cm) and their behavior while freely exploring was monitored for 10 min using a video camera mounted on the ceiling above the center of the arena. For analysis, the floor was divided into three virtual concentric parts, with a center zone in the middle of the arena (20 cm diameter), an interior zone (60-cm diameter), and an exterior zone made up of the remaining area along the sidewalls. Different parameters were evaluated with the video tracking system: distance moved (centimeters) and time spent (seconds) in each zone. Immediately after the open field test, rats were submitted to the novel object reactivity (NOR) test. For this purpose, a small, white plastic bottle was placed into the center of the open field while the rat was inside. Rats were then given 5 min to freely explore the novel object. The time spent exploring (touching) the novel object was recorded manually from the video recordings. Moreover, different parameters were evaluated with the video tracking system: time spent (seconds) in the center (where the novel object was placed) and the periphery of the compartment, number and latency of entries to the center, total distance moved (centimeters) in the center and in the whole compartment.

### Water Maze

In order to test spatial learning and memory, a round black Plexiglas tank with a diameter of 2 meters and a height of 45 cm was used. The pool was filled with water each day and the temperature was maintained at 25°C ± 1°C during the experiment. A circular platform was submerged 1.3 cm below the water surface. The water maze was surrounded by clearly discernible visual cues to facilitate spatial orientation during the training phase. The experiment was divided in two phases: training and probe trial. The training phase lasted from Day1 to Day 3 and it involved 4 × 90-second trials/day/rat with a 30 seconds inter-trial interval. The platform remained constantly at the quadrant assigned as the target quadrant. The starting point for each trial was pseudo-randomly chosen. In order to assess the spatial memory of the animals, a probe trial was performed 24 hours after the last training session (Day 4). During this phase, the platform was removed and rats were allowed to swim freely for 90 seconds. The distance that the animals swam to find the platform was used as an indication of learning. The percentage of the time spent in the quadrant that contained the platform during training (target quadrant) versus the adjacent quadrant was used as an index of spatial memory.

### Corticosterone Responsiveness

Individual responsiveness to PPS was evaluated in the STRESS group by measurement of plasmatic corticosterone (CORT) levels during the first and last day of PPS exposure. Blood samples were obtained by tail-nick protocol (100 μl for peripubertal animals) within 2 min while gently holding the animals with a cloth and, then, animals were returned to their home cage. The tail-nick procedure allows for the collection of blood samples at different time points from the same animal (Márquez et al., 2004), which enables the study of hormonal dynamics. Samples were obtained in basal conditions, immediately following the termination of the elevated platform stress and 30 and 60 min after the elevated platform stress. Based on these CORT measurements, two adaptation indices were then calculated: time 0 (t0) and recovery 60 (r60). The t0 index reflected the change of the CORT response, immediately after exposure to the stressor, between the last (P42) and the first (P28) day of the protocol (CORT P42 immediately after stress * 100/ CORT P28 immediately after stress), and thus, expressed a proxy for the adaptation of the initial response to the stressor after subchronic stress. In a similar way, the recovery 60 (r60) index, was calculated to assess recovery adaptation to basal corticosterone levels after exposure to stress (CORT P42 60 min after termination of stress exposure * 100/ CORT P28 60 min after termination of stress exposure). Two animals, one from the adolescent group and one from the adulthood group were excluded from all analyses as the values for these variables were exceeding 3 Standard Deviations from the mean.

Later in life, the long-term effects of PPS on corticosterone reactivity to a mild stressor were evaluated in independent groups of CTRL and STRESS animals at either adolescence or adulthood period. Immediately after 30 min exposure to a novel environment (a circular corridor made of plastic; 35 cm high, 25 cm diameter) blood samples were obtained by tail-nick (250 μl). Two additional blood samples were obtained during the recovery period, 30 and 60 min after the end of circular corridor exposure. Baseline samples were collected in a previous day, in order not to interfere with behavior. Animals from the same home-cage were simultaneously tested in adjacent containers. The containers were cleaned with 1% acetic acid and dried properly before placing the animals.

Blood samples were collected into ice-cold heparin capillary tubes (Sarsted, Switzerland) and kept at 4 degrees during the experiment. Plasma was obtained after blood centrifugation at 10,000 rpm for 25 min and stored at –20°C until analyses. Plasma corticosterone levels were measured by enzymatic immunoassay kit (Correlate-EIA from Assay Designs Inc., USA) according to supplier’s recommendations. The area under the curve of the corticosterone levels was calculated using GraphPad Prism (version 7), which computes the area under the curve using the trapezoid rule.

### PSA-NCAM Immunohistochemistry

For the PSA-NCAM immunohistochemistry experiment rats were anesthetized with a lethal dose of pentobarbital (Esconarkon, Streuli Pharma AG, 150 mg/kg body weight, solution provided by the EPFL veterinarian) and perfused via the ascending aorta with ice-cold 0.9% saline, followed by 4% paraformaldehyde in phosphate-buffered saline (pH=7.5). After perfusion-fixation, the brains were removed from the skull, post-fixed in the same solution for four hours, and stored in 4°C PBS until further processing. Subseries of 50 μm thick sections from each group of animals were processed free floating for immunohistochemistry using the avidin-biotin-peroxidase (ABC) method (Hsu et al, 1981). Sections were incubated with 10% H_2_O_2_ in phosphate buffered saline (PBS) for 10 minutes to block endogenous peroxidase activity. They were then treated for 1 hour with 10% normal donkey serum (NDS) (Jackson ImmunoResearch Laboratories) in PBS with 0.2% Triton-X100 (Sigma-Aldrich) and incubated for 60 hours at 4°C in the primary antibody anti-PSA-NCAM, generated in mouse, (DSHB, 1:1500) with PBS containing 0.2% Triton-X-100. Then, sections were incubated for 2 hour at RT with the biotinilated secondary antibody: donkey anti-mouse IgG (Jackson ImmunoResearch Laboratories, 1:200), followed by an avidin-biotin-peroxidase complex (ABC; Vector Laboratories) for 1 hour in PBS. Color development was achieved by incubating with 3,3’-diaminobenzidine tetrahydrochloride (DAB; Sigma-Aldrich) and 0.033% H_2_O_2_ for 4 minutes. Finally, sections were mounted on slides, dried for one day at room temperature, dehydrated with ascending alcohols and rinsed in xylene. Sections were coverslipped using Eukitt mounting medium (PANREAC). All sections passed through all procedures simultaneously in order to minimize any difference from the immunohistochemical staining itself. To avoid any bias in the analysis, all slides were coded prior to analysis and remained so until the experiment was completed. Sections were examined with an Olympus CX41 microscope under bright-field illumination, homogeneously illuminated and digitalized using a CCD camera. Photographs of the different areas were taken at 20Å∼ magnification. Grey levels were converted to optical densities (OD) using Image J software (NIH). Means were determined for each experimental group and data were analyzed with appropriate statistical tests.

### Statistical Analyses

During behavioral testing animals were tracked automatically with EthoVision 3.0/3.1 (Noldus, Wageningen, the Netherlands). The results were analyzed using the SPSS 17 statistical package and the graphs and correlation matrices were made using GraphPad Prism 7. The data were analyzed with analysis of variance (ANOVA) with repeated measures, Student’s t – tests or paired samples t-tests as considered appropriate. The data was checked for distribution with the Shapiro-Wilk test and when the normality was violated, non-parametric tests were applied (i.e. Mann-Whitney and Wilcoxon signed rank tests). All t-tests and paired samples t-tests were performed two-tailed, with the exception of the Mann-Whitney for Fig.3I and 3J, where we specifically hypothesized a blunted CORT response (i.e., one-tailed prediction) extrapolating from previous findings (Veenit et al., 2013). Regarding the repeated measures ANOVA, when Mauchly’s test of sphericity was significant, thus sphericity could not be assumed, the Greenhouse-Geisser correction was used and reported. All results represent the mean + the standard error of the mean (SEM) and the significance was set at p < 0.05.

## Results

### Marked individual differences in the corticosterone adaptation to repeated stressor exposure during the peripubertal period

In order to determine whether adaptation to the PPS protocol (peak vs recovery) could predict long term reprogramming effects of stress, we first characterized the corticosterone (CORT) response dynamics during stress exposure. Rats were exposed to threatening challenges (i.e., elevated platform, predator odor) at scattered days (i.e., P28, P29, P30, P34, P36, P40 and P42) within the peripubertal period (Fig. 2A) and blood samples collected following exposure to the same stressor, elevated platform, on the first (P28) and last (P42) days of the protocol, and at four time points: immediately before the stressor (basal), immediately after the elevated platform exposure (stress; t0) and in order to assess the recovery of the response, 30 and 60 minutes following the stressor (rec30 and rec60 respectively). In both days, exposure to the elevated platform induced a robust corticosterone release that was recovered to basal levels 1 hour after stress termination (Fig 2B; left). A general habituation of the CORT response from P28 to P42 was observed, with CORT levels being reduced at P42, as indicated by a decreased AUC measurement (Fig. 2B; right: *t* (32) = 9.398, *p* < 0.001). Importantly, inspection of these results indicates that rats displayed marked individual differences in their corticosterone adaptation to peripubertal stress, both as in their peak responses (Fig. 2C) and in the rec 60 time-point (Fig. 2D). Thus, while the majority of animals showed a decreased CORT response at P42 with varying levels of intensity, suggesting a good degree of adaptation, a subset of rats did not adapt at all (Fig. 2C-D). As expected, the same pattern of CORT response was obtained when animals - ascribed to the two testing conditions later in life - were analyzed separately for validation purposes (Fig. S1 A-D; adulthood: A – left; Wilcoxon signed rank paired test – basal: *p* < 0.001, stress: *p* = 0.001, rec 30: *p* < 0.001, rec 60: p < 0.001, A – right; *t* (16) = 7.817, *p* < 0.001), adolescence: C – left; Wilcoxon signed rank paired test – basal: *p* < 0.001, stress: *p* = 0.001, rec 30: *p* = 0.004, rec 60: *p* = 0.002, C – right; *t* (15) = 5.830, *p* < 0.001, B and D; animals plotted individually for t0 and rec60 time points for adulthood and adolescence respectively. Animals showing low adaptation can be observed in both age groups).

### Peripubertal stress leads to delayed programming effects on anxiety-like behavior

Before assessing for potential programming effects of peripubertal stress in spatial learning, we tested animals for their locomotor and exploratory behaviors in the Open Field and Novel Object tests, as behavioral changes in these tests may help interpreting potential differences in the water maze. When tested at adulthood, STRESS rats showed a decrease in the time spent in the center of the Open Field (Fig. 3A; Mann-Whitney – *p* = 0.014), no differences in total distance moved (Figure 2B; *t* (31) = −0.940, *p* = 0.354), but an increase in self-grooming behavior (Fig. 3C; Mann-Whitney test, *p* = 0.007). In addition, STRESS rats showed a trend towards increased time exploring and touching the object in the novel object test (Fig. 3D; t (31) = −1.769, *p* = 0.087). Altogether, these results indicate a phenotype characterized by increased anxiety-like behaviors with no change in locomotion.

**Figure 3.**
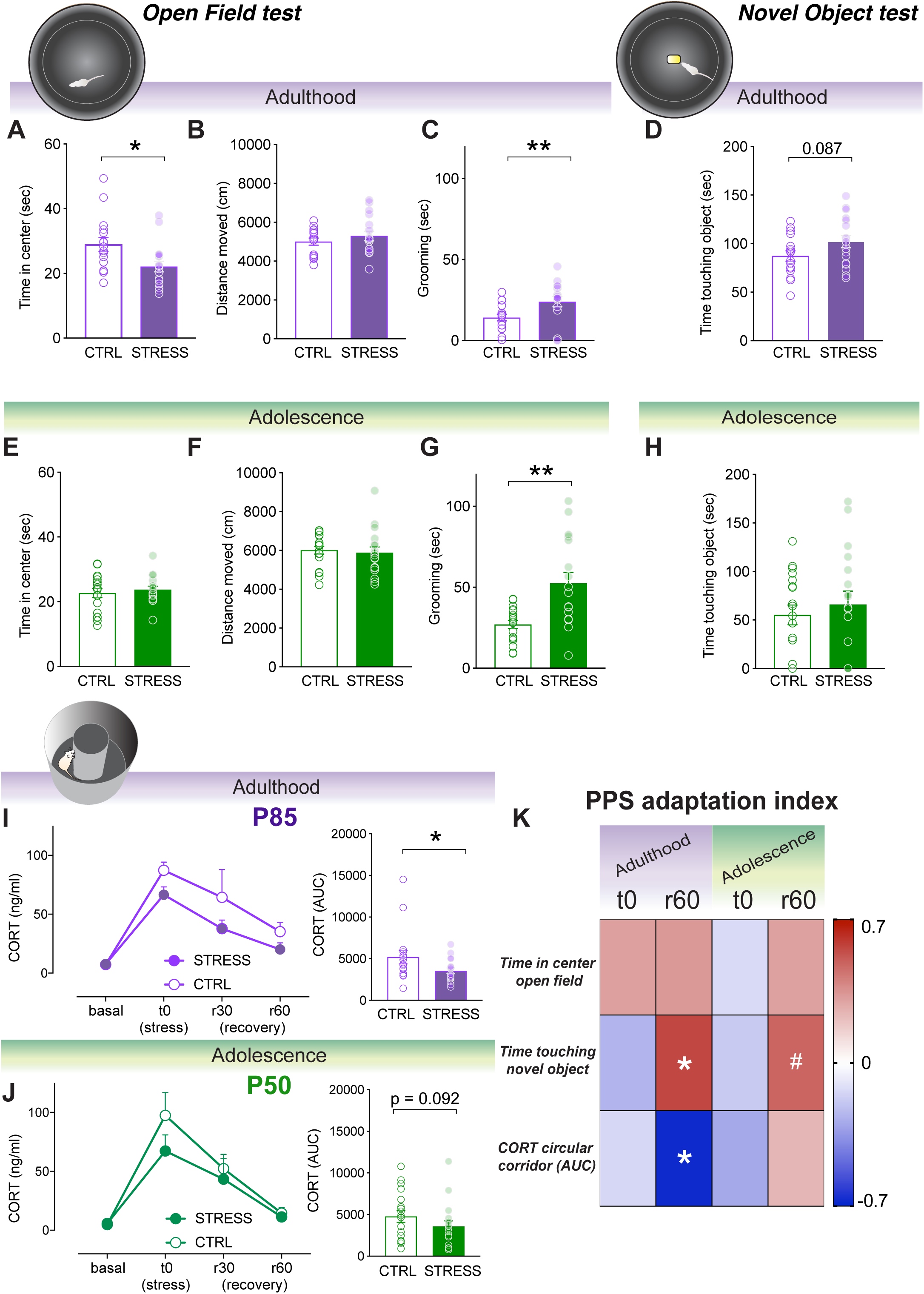
Open Field, Novel Object tests and corticosterone reactivity to circular corridor for adult and adolescent CTRL and STRESS rats. Adult STRESS rats spent less time in the central area of the Open field test (**A**) but no differences in general locomotor activity were observed (**B**). Moreover, self-grooming levels observed in the open field were also higher in STRESS animals (**C**). Regarding exploration, STRESS animals tended to explore more the object in the novel object test (**D**). None of these differences were observed in adolescence (**E, F, H**) except for the increase of grooming levels during open field exposure (**G**). CORT response was also examined upon exposure to a novel environment (i.e. circular corridor) in either CTRL or STRESS rats in adulthood. **I**. STRESS rats exhibited lower CORT response upon exposure to a novel environment (i.e. circular corridor) compared to CTRL animals. Temporal corticosterone dynamics are presented in the left, and AUC is shown in the right. **J**. The same tendency was observed in adolescence; however, it did not reach statistical significance. K. Correlation matrix between the adaptation index after peripubertal stress considering peak (t0) or recovery times (r60) and main behavioral and endocrine measurements. Animals that adapted the less during the recovery to peripubertal stress were the more impaired in the novel object test and with a more marked blunted CORT response to stress during adulthood. Results are expressed as the mean ± S.E.M. *p<0.05, **p<0.01, # p<0.10

In order to ascertaining whether these behavioral changes emerged at adulthood or were already present at earlier time points, we tested a second cohort of animals during adolescence (P55+; Figure 1). At this time point, STRESS animals did not show changes in the time spent in the center (Fig. 3E; *t* (33) = −0.606, *p* = 0.548) or distance moved (Figure 2F; *t* (33) = 0.366. *p* = 0.717) in the open field. However, as when tested at adulthood, STRESS animals tested at adolescence showed increased self-grooming behavior (Fig. 3G; *t* (20.434) = −3.584, *p* = 0.002). In the novel object test (Fig. 3D; t (31) = −1.769, *p* = 0.087), they did not differ from CTRL in time exploring the object (Fig.3H; *t* (33) = −0.626, *p* = 0.535).

Therefore, these data indicate an interesting age-dependent effect on exploratory behaviors. Specifically, long-term programming effects of peripubertal stress on anxiety-like behaviors are observed at adulthood, and in a much lesser extent (i.e., self-grooming) at adolescence. Locomotion is not changed at any of the testing times.

### Peripubertal stress induces CORT hypo-reactivity in adulthood

We then sought to ascertaining if exposure to PPS would affect corticosterone reactivity to challenges later in life. Indeed, adult STRESS rats showed blunted CORT reactivity (Fig. 3I - left: main effect of time: *F* (1.4, 43.68) = 11.434, *p* < 0.001, main effect of stress: *F* (1, 31) = 3.861, *p* = 0.058, stress x time interaction: *F* (1.4, 43.68) = 0.152, *p* = 0.783, 3I - right: Mann-Whitney one tail test, *p* = 0.0265) following exposure to a novel environment (i.e., circular corridor, devoid of the anxiogenic center of the arena). This effect was particularly obvious at adulthood, as a mild reduction in CORT activation observed when STRESS animals were tested during adolescence was not significant (Fig. 3J - left; main effect of time: *F* (1.5, 49.05) = 14. 565, *p* < 0.001, main effect of stress: *F* (1, 33) = 1.547, *p* = 0.222, stress x time interaction: *F* (1.5, 49.05) = 0.615, *p* = 0.498, 3J - right: Mann-Whitney one tail test, *p* = 0.092).

Then, we aimed to understand whether the degree to which animals adapt their corticosterone responses to repeated stressors during the peripubertal period (i.e., from P28 to P42) relates to subsequent behavioral and/or hormonal responses. To this end, we first computed two adaptation indices for time 0 (t0) and recovery 60 (r60) (see Methods for details). Then, we calculated correlations between these indices and key behavioral parameters and corticosterone reactivity (AUC) to emotional challenges (i.e., the tests reported above). As shown in Fig. 3K, it was specifically the adaptation of CORT recovery (rec60 index) during peripubertal stress that correlated with both, time touching the object in the Novel object test (Fig. 3K; r = 0.558, *p* = 0.020) and, negatively, with corticosterone reactivity (r = −0.670, *p* = 0.003) in animals tested at adulthood. Thus, the lesser the adaptation of corticosterone recovery during PPS stress exposure, the lower the time exploring the novel object, and the lower the CORT responsiveness a mild stressor (*i*.*e*. novel environment). A similar trend, although not significant, was observed for the correlation between time touching the novel object and rec60 index for the data from adolescence testing (Fig. 3K; r = 0.481, *p* = 0.059). Strikingly, no correlation was observed between the studied parameters and the PPS CORT adaptation index for t0 (i.e, peak CORT stress responses).

### Peripubertal stress leads to delayed programming effects on spatial learning

We then addressed our main question; whether peripubertal stress can have delayed effects on spatial learning and memory, and to what extent any observed effect would be related to the degree of CORT adaptation to repeated stressor exposure during peripuberty. To this end, animals were trained and tested to find a hidden platform in the water maze. Adult STRESS rats showed increased total distance swam to find the hidden platform on day 2 (Fig. 4A; left - *day 1*-main effect of time: *F* (3, 48) = 4.192, *p* = 0.010, main effect of stress: *F* (1, 16) = 0.295, *p* = 0.595, time x stress interaction: *F* (3, 48) = 0.109, *p* = 0.955, *day 2* - main effect of time: *F* (3, 48) = 3.231, *p* = 0.030, main effect of stress: *F* (1, 16) = 5.110, *p* = 0.038, time x stress interaction: *F* (3, 48) = 0.085, *p* = 0.968, *day 3* - main effect of time: *F* (3, 48) = 2.182, *p* = 0.102, main effect of stress: *F* (1, 16) = 2.923, *p* = 0.107, time x stress interaction: *F* (3, 48) = 0.797, *p* = 0.502)) and increased distance moved when the average performance per session was considered (Fig. 4B; middle - main effect of time: *F* (2, 32) = 17.083, *p* < 0.001, main effect of stress: *F* (1, 16) = 5.713, *p* = 0.029, time x stress interaction: *F* (2, 32) = 1.049, *p* = 0.362). No differences between STRESS and control rats were observed during the probe trial (Fig. 4B; right - adjacent: *t* (16) = - 0.068, *p* = 0.947, target: *t* (16) = 0.068, *p* = 0.946).

**Figure 4.**
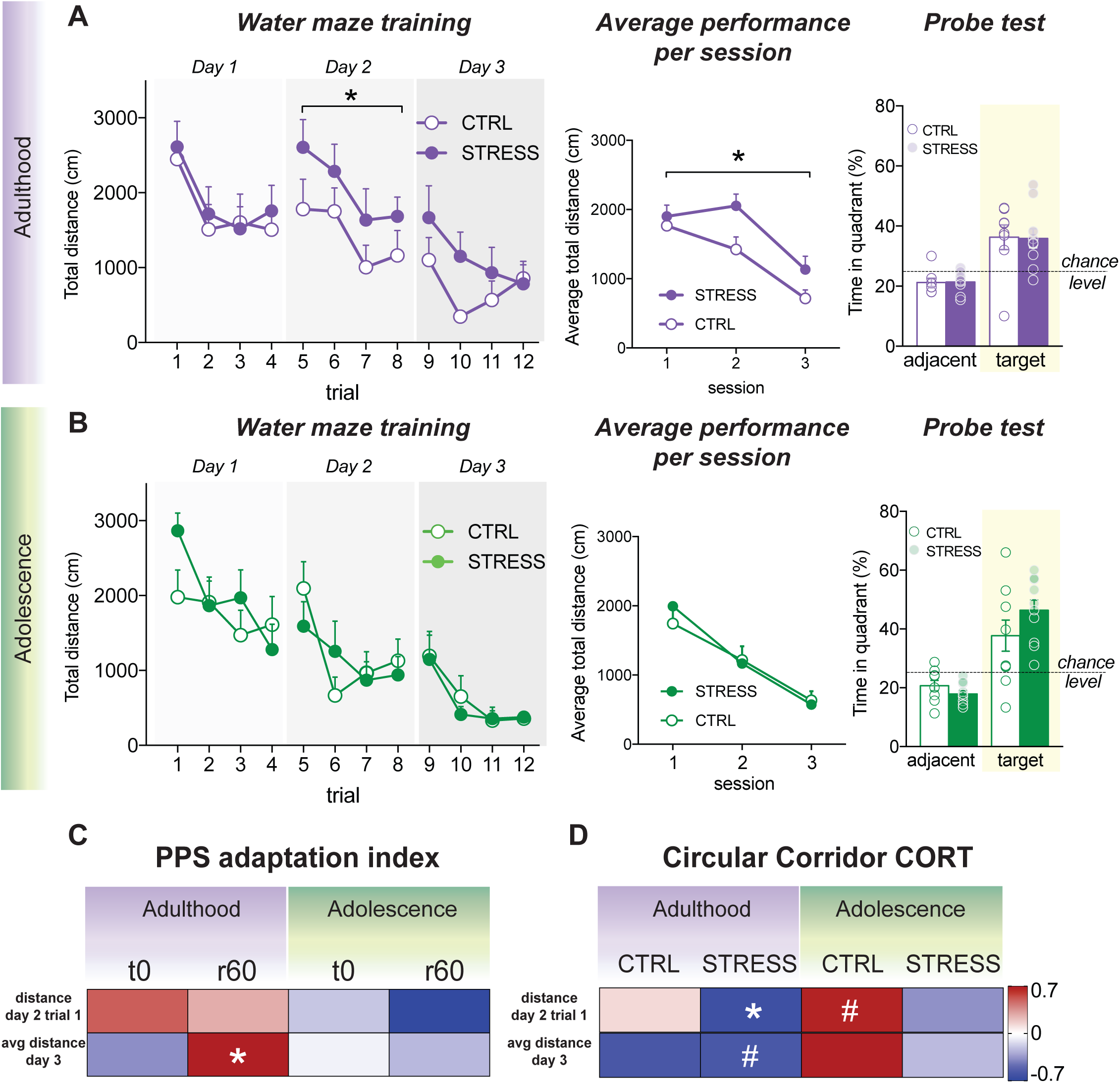
Water maze training and probe trial for adult and adolescent rats, as well as correlations of water maze parameters with CORT. **A**. Left: Adult STRESS rats traveled more distance before finding the platform on the second day of training, compared to CTRL rats. Middle: Compared to CTRL, STRESS rats tested during adulthood showed increased distance in the water maze when the average performance per session was considered. Right: Both CTRL and STRESS adolescent groups exhibited intact memory of the position of the platform during the probe trial, when the platform was absent. No differences were found between the groups. **B**. Left: No differences were observed between adolescent CTRL and STRESS rats during training in the water maze. Middle: No differences between the groups were observed for the average performance per session. Right: Both CTRL and STRESS adolescent groups exhibited intact memory of the position of the platform during the probe trial, when the platform was absent. No differences were found between the groups. **C**. Correlation matrix showing correlation coefficients between peak (t0) and recovery (r60) adaptation indexes to peripubertal stress and performance in a in the first long term memory test (trial 1 of the second day of testing) and in the last day of training. Those animals that adapted the less to peripubertal stress (r60) were the more affected while adults and performed the worse in the Morris water maze. **D**. Correlation matrix between circular corridor CORT response and performance in the water maze, indicating that only in STRESS animals, a more blunted CORT response to the novel environment (lower CORT levels) correlated with impairments in the Morris water maze. Results are expressed as the mean ± S.E.M. Correlation coefficients (r values) in correlation matrix are color-coded. *p<0.05, # p<0.10

In order to inquiry whether the observed PPS stress effects on spatial learning at adulthood were protracted or delayed, we tested the second cohort of animals in the water maze during adolescence. However, at this time point, no effect of PPS stress was observed (Fig 4B; left – no effect in distance moved: *day 1*-main effect of time: *F* (3, 54) = 3.306, *p* = 0.027, main effect of stress: *F* (1, 18) = 0.832, *p* = 0.374, time x stress interaction: *F* (3, 54) = 1.455, *p* = 0.237, *day 2* - main effect of time: *F* (3, 54) = 5.379, *p* = 0.003, main effect of stress: *F* (1, 18) = 0.032, *p* = 0.860, time x stress interaction: *F* (3, 54) = 1.476, *p* = 0.231, *day 3* - main effect of time: *F* (1.76, 31.67) = 8.329, *p* = 0.002, main effect of stress: *F* (1, 18) = 0.108, *p* = 0.746, time x stress interaction: F (1.76, 31.67) = 0.217, p = 0.778) nor regarding the average performance per session (Fig. 4B; middle - main effect of time: F (2, 36) = 44.392, p < 0.001, main effect of stress: *F* (1, 18) = 0.058, *p* = 0.812, time x stress interaction: *F* (2, 36) = 0.871, *p* = 0.427), nor in the probe test (Fig. 4B; right - adjacent: *t* (18) = 1.360, *p* = 0.191, target: *t* (18) = −1.361, *p* = 0.190).

We then inquired whether individual differences in CORT adaptation during PPS exposure (i.e., t0 and rec60 indices) related to differences in key parameters of water maze performance. To this end, we selected average performance (i.e., distance to the platform) on the last training day, as an index for the maximal acquisition level obtained) and distance to find the platform on the first trial of day 2, as a first long-term memory index. As shown in Fig. 4C, rec60 was again the parameter that showed a positive correlation with day 3 performance (Fig. 4C; r = 0.648, *p* = 0.043); i.e., the lesser the adaptation of CORT recovery following peripubertal stressors, the worse the maximal training performance in the water maze. This was not observed when the animals were tested in adolescence (Fig. 4C). Moreover, no correlations were found between water maze performance and PPS peak adaptation index (t0 index) at any of the age groups (Fig 4C).

Furthermore, in order to better understand possible links between CORT responsiveness during the testing period and variation in spatial learning performance, we examined the relationship between CORT response upon exposure to the circular corridor (see Figs 3I, J) and water maze parameters. Interestingly, CORT reactivity to the circular corridor correlated with the first long-term memory test (i.e., distance to find the platform on the first trial of day 2) in STRESS animals tested at adulthood; i.e., the lower the CORT the poorer their performance in the water maze (Fig. 4D; r = −0.641, *p* = 0.046). In other words, those animals that displayed more blunted corticosterone response during adulthood after a novelty challenge as a consequence of peripubertal stress exposure were the ones showing worst long-term memory in the water maze. A similar trend was observed for STRESS animals’ CORT responsiveness at adulthood and average performance on the last training day (Fig. 4D; r = −0.581, *p* = 0.078).

Altogether, these results suggest that peripubertal stress has delayed detrimental effects on spatial learning that become evident when the assessment happens during adulthood, and that those animals that show impaired adaptation in CORT recovery to repeated stressors exposure perform poorer in a spatial learning task.

### Peripubertal stress leads to changes in PSA-NCAM in the dentate gyrus

In order to gain insight into key plasticity molecules, related to learning and memory, that could be affected by peripubertal stress, two further cohorts of rats were exposed to peripubertal stress and studied at each age group (i.e., CTRL and STRESS; Adulthood and Adolescence) and assessed for the expression of PSA-NCAM in the DG of the hippocampus (Fig. 5). As shown in Fig. 5A, there was an increase of PSA-NCAM in adult rats stressed during peripuberty (Fig. 5A – left; *t* (12) = −3.675, *p* = 0.003). However, no significant differences were apparent in PSA-NCAM expression when the rats were assessed during adolescence (Fig. 5A – right; *t* (13) = −1.458, *p* = 0.169). In order to study specificity of our findings, we quantified PSA-NCAM expression in the medial amygdala. However, no differences were found between STRESS and CTRL rats regardless of the developmental age, suggesting that the PSA-NCAM alterations observed in the dentate gyrus at adulthood were not only age-dependent, but also brain region-specific (Fig. 5C – left – adulthood; *t* (8.04) = −1.059, *p* = 0.321, right – adolescence; *t* (13) = −0.291, *p* = 0.776). Interestingly, there was a trend for those animals whose CORT responsiveness would adapt suboptimally at r60 during peripubertal stress to display higher DG PSA levels in adulthood (r = 0.735 *p* = 0.096) (Fig 5D). No correlations of CORT responsiveness were observed with the MeA for any of the age groups or time points.

**Figure 5.**
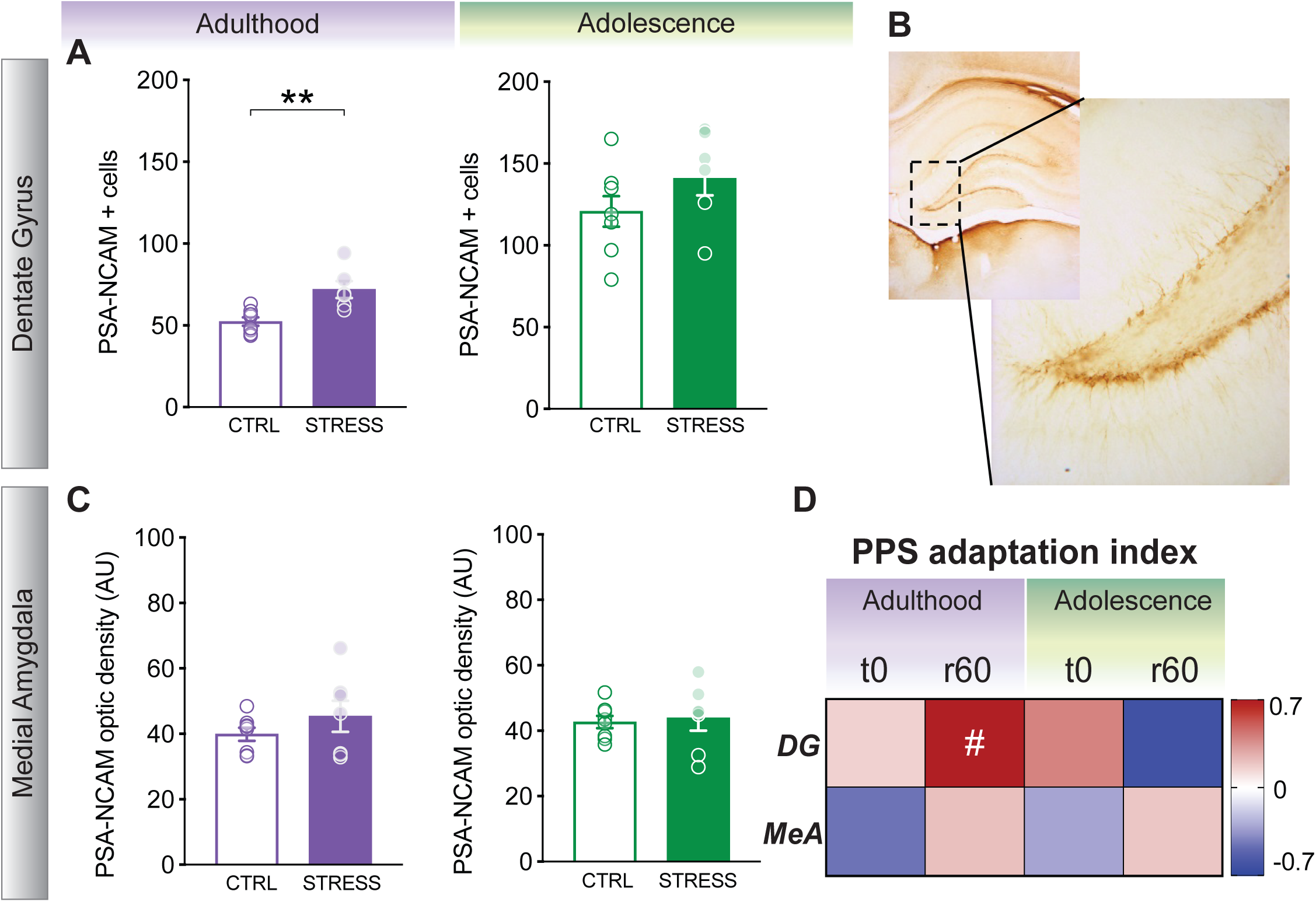
PSA-NCAM levels for adult and adolescent CTRL and STRESS rats in the Dentate Gyrus (DG) and Medial Amygdala (MeA). **A**. Left: Adult STRESS rats showed increased number of PSA-NCAM positive cells compared to adult CTRL rats. Right: These differences were not observed when assessed during adolescence. **B**. Photomicrograph of the DG area where PSA-NCAM positive cells were quantified. **C**. No difference was observed between CTRL and STRESS tested at adulthood (left) nor adolescence (right) in PSA-NCAM optic density in the MeA. **D**. Correlation matrix showing correlation coefficients between peak (t0) and recovery (r60) adaptation indexes to peripubertal stress and PSA-NCAM levels in the dentate gyrus and medial amygdala. Those animals that adapted the less to peripubertal stress (r60 index) were the ones that tended to display higher levels of DG PSA-NCAM during adulthood. Results are expressed as the mean ± S.E.M. Correlation coefficients (r values) in correlation matrix are color-coded. **p<0.01, # p<0.10

## Discussion

Here, we show that exposure to stressors across the peripubertal period in rats leads to cognitive, behavioral and endocrine changes at adulthood. Specifically, peripubertal stress led to impaired spatial learning, increased anxiety-like behavior, and blunted corticosterone responsiveness to novelty challenges. These effects are delayed in nature, as they were not displayed by animals tested during adolescence (i.e., shortly after peripubertal stress exposure). Strikingly, individual differences in the degree of adaptation of the recovery -and not the peak-of the corticosterone response to stressor exposure (i.e., plasma levels at 60 min post-stressor) across the peripubertal stress period (i.e., from P28 to P42) predicted the level of spatial orientation in the water maze completed on the last training day, as well as the exploratory behavior shown by animals at adulthood. In addition, this corticosterone stress adaption recovery index (rec60) was inversely related to the corticosterone responsiveness to novelty at adulthood. These findings contribute to further our understanding on the link between HPA axis adaptation to early life stressors at the important transitional period of puberty and the long-term programming of behavior and cognition.

Thus, a main finding of our study is the identification of peripuberty as a stress-sensitivity period for the modulation of adult spatial learning abilities. Previous studies had underscored the early postnatal period as a time-window in which stress exposure makes individuals particularly prone to show spatial learning impairments at long-term life stages (Brunson et al., 2005) (Brunson et al., 2005; Oomen et al., 2010). However, previous studies comprising stressor exposure across several weeks from juvenility to adulthood in which spatial learning and memory impairments were reported (Isgor et al., 2004; Sterlemann et al., 2010) did not allow disentangling the putative impact of peripubertal stress *per se*. In addition, we show here that the impact is not immediate (i.e., not shown during adolescence) but, similarly to the report by Isgor et al. (2004), it only emerges when testing takes place several weeks after the end of the stress protocol, at adulthood. Further evidence for this delayed phenomenon stems from studies in rats involving prepubertal stress (i.e., from P28 to P30) and showing impaired water maze at adulthood only following a second stressful challenge at adulthood that, on its own, does not affect spatial learning performance (Avital and Richter-Levin, 2005). In this connection, we previously reported that the same peripubertal stress protocol as the one applied here leads as well to attention deficits in adulthood (Tzanoulinou et al., 2016), but whether these deficits are observed already during adolescence remains to be tested. Altogether, these findings support the view that the long-term cognitive impact of peripubertal stress requires an incubation period during which stress-targeted mechanisms interact with ongoing maturational and neurodevelopmental trajectories to produce phenotypic changes at later life stages.

Spatial learning highly depends on the functioning of the hippocampus (Moser et al., 1995), a brain region that undergoes profound structural and functional changes in adolescence (McCormick and Mathews, 2010). Interestingly, efficient spatial orientation strategies in the water maze task appear around P42 (Schenk, 1985), coinciding with the last day of our peripubertal stress protocol. Our own data on the expression levels of the plasticity molecule PSA-NCAM in the dentate gyrus show a down-regulation from adolescence to adulthood; this age-dependent regulation was not observed in the medial amygdala, a brain region not involved in spatial learning. Thus, our data agrees with an age-dependent pattern of PSA-NCAM down-regulation taking place in the brain during the postnatal period (Angata and Fukuda, 2003; Rutishauser, 2008) and remaining present later in life in brain areas that maintain neurogenic potential or heightened plasticity, such as the hippocampus (Angata and Fukuda, 2003), where it has been causally involved in memory consolidation (Doyle et al., 1992; López-Fernández et al., 2007; Sandi et al., 2003; Venero et al., 2006). Importantly, we found that peripubertal stress leads to increased PSA-NCAM levels specifically in the dentate gyrus, that was particularly evident in the group of animals examined at adulthood, in agreement with similar findings following exposure to pre-pubertal/juvenile stress (Tsoory et al., 2008). These results reflect the effectiveness of peripubertal stress to disrupt the maturation of the hippocampal learning system. PSA can promote neuronal and synaptic plasticity through mechanisms involving its de-adhesive properties, as well as by interacting with extracellular matrix molecules and glutamate receptors (Varbanov and Dityatev, 2017)). However, while PSA-NCAM expression in the dentate gyrus transiently increases around 12 h following training in the water maze (Murphy et al., 1996; Sandi et al., 2003), this increase is only observed in bad -but not good-learners, that require increased effort to complete the task (Sandi et al., 2004), and it decays as animals progressively master the task (Murphy et al., 1996). Moreover, chronic stress at adulthood leads to increased hippocampal PSA-NCAM expression (Pham et al., 2003; Sandi et al., 2001) and impairs spatial learning in the water maze (Sandi, 2004; Venero et al., 2002). Therefore, the facilitation of learning and plasticity processes by PSA-NCAM seems to require an activity-dependent process triggering a transient increase in its expression. Heightened basal elevation of PSA-NCAM appears to be deleterious to information processing, which aligns with our findings in the current study. Furthermore, we should also note that we found a trend for dentate gyrus PSA-NCAM levels to correlate with the adaptation of the corticosterone recovery levels across the peripubertal stress protocol (rec60 index; i.e., the lower the adaptation, the higher PSA-NCAM levels). Although until further replication these findings should be taken with caution given the reduced sample size, they point towards a potential role of glucocorticoids on the regulation of hippocampal PSA-NCAM expression by peripubertal stress. Indeed, a complex regulation of hippocampal PSA-NCAM by glucocorticoids has been revealed (Nacher et al., 2004; Rodríguez et al., 1998) Rodriguez et al., 1998; Nacher et al., 2004), involving, in particular, glucocorticoid receptor actions (Montaron et al., 2003).

Importantly, the peripubertal period entails a transition in HPA responsivity to stressors at both, peak and recovery phases (McCormick et al., 2017). Strikingly, we found that individual differences in the spatial orientation levels achieved in the last training day were also related to the peripubertal rec60 corticosterone adaptation index. Thus, those animals that showed a poorer adaptation of the corticosterone stress recovery at puberty were the ones that attained poorer performance levels. Importantly, as hypothesized, it was the corticosterone recovery, and not the peak, index that related to water maze performance. The ability to down-regulate the HPA axis response to stress (and thus, corticosterone levels) following stress exposure through negative feedback is essential to protect the organism from maladaptive overactivation (Karatsoreos and McEwen, 2011) and also important for optimal secretion of corticosterone in basal (unstressed) conditions (Gjerstad et al., 2018). Therefore, our rec60 index seems to have captured individuals’ ability to adapt to repeated life stressors and serves as a predictive index of adult life cognitive, behavioral, and endocrine disturbances. These findings align well with the important role of the hippocampus in providing negative feedback to the HPA axis (Herman and Mueller, 2006; Jacobson and Sapolsky, 1991; Kovács and Makara, 1988) and the involvement of hippocampal glucocorticoid receptors in HPA axis negative feedback (de Kloet Frontiers in neuroendocrinology Print1991, n.d.; Reul et al., 1990).

In addition, the first index of long-term memory performance (i.e., distance moved to find the platform on the first trial of training day 2) was related to corticosterone reactivity at adulthood. Specifically, animals that showed poorer retention levels on the first trial following training day 1 were the ones that showed blunted corticosterone reactivity when exposed as adults to a novelty challenge. These observations align well with the well-known contribution of training-triggered corticosterone levels for memory function in general (de Kloet et al., 2018; de Quervain et al., 2017; Sandi, 2011)) and, specifically, for the consolidation of spatial information (Akirav et al., 2004; Conboy et al., 2010; Huzard et al., 2020; Quirarte et al., 1997; Sandi et al., 1997).

The incubation period reported here for spatial learning effects of peripubertal stress to emerge at adulthood appears to be specific for the cognitive domain. Indeed, a different process seems to be engaged in the development of anxiety-like behaviors. While, as in previous studies (Cordero et al., 2016; Tzanoulinou et al., 2014a)), we observe here increased anxiety-like behavior when peripubertally stressed rats were tested at adulthood, decreased anxiety-like behaviors were reported when tested during late adolescence (Toledo-Rodriguez and Sandi, 2011). Furthermore, in the social domain, peripubertal stress leads to increased adult aggression (Marquez et al., 2013) in a protracted manner, as rats that showed aberrant play fighting during adolescence were those that developed a more aggressive phenotype at adulthood (Papilloud et al., 2018). In addition, and in line with its physiological contribution to deal with immediate challenges (de Kloet et al., 2008; Myers et al., 2014), it is the magnitude of adaptation of the peak corticosterone response to peripubertal stress that predicts alterations in emotional and social behaviors (Papilloud et al., 2018; Walker et al., 2018; 2017). Along the same lines, we demonstrated here, that the ability to adapt during recovery periods, i.e. once stress exposure has ended, is a key predicting factor for special learning impairments.

In summary, our study identifies the peripubertal period as a time-window at which stress can lead to long-term changes in HPA axis reactivity that are related to difficulties in spatial learning abilities later in life. These findings pave the way for further studies to identify mechanisms of both vulnerability and resilience to early trauma. Furthermore, our data suggest that the reprograming effects of early stress might need a period of incubation which could be compensated in young and more plastic brains, but would fail to adapt during adulthood. Accordingly, following early detection of stress-vulnerable individuals, there may be a window of opportunity for therapeutic approaches to act during adolescence deflecting the course trajectory towards psychopathology and cognitive impairments.

## Acknowledgements

We thank Angélique Voucher, Coralie Siegmund, Marjorie Clerc and Aliénor Sonnay for excellent technical assistance.

## Supplementary Figures

**Supplementary Figure 1.**
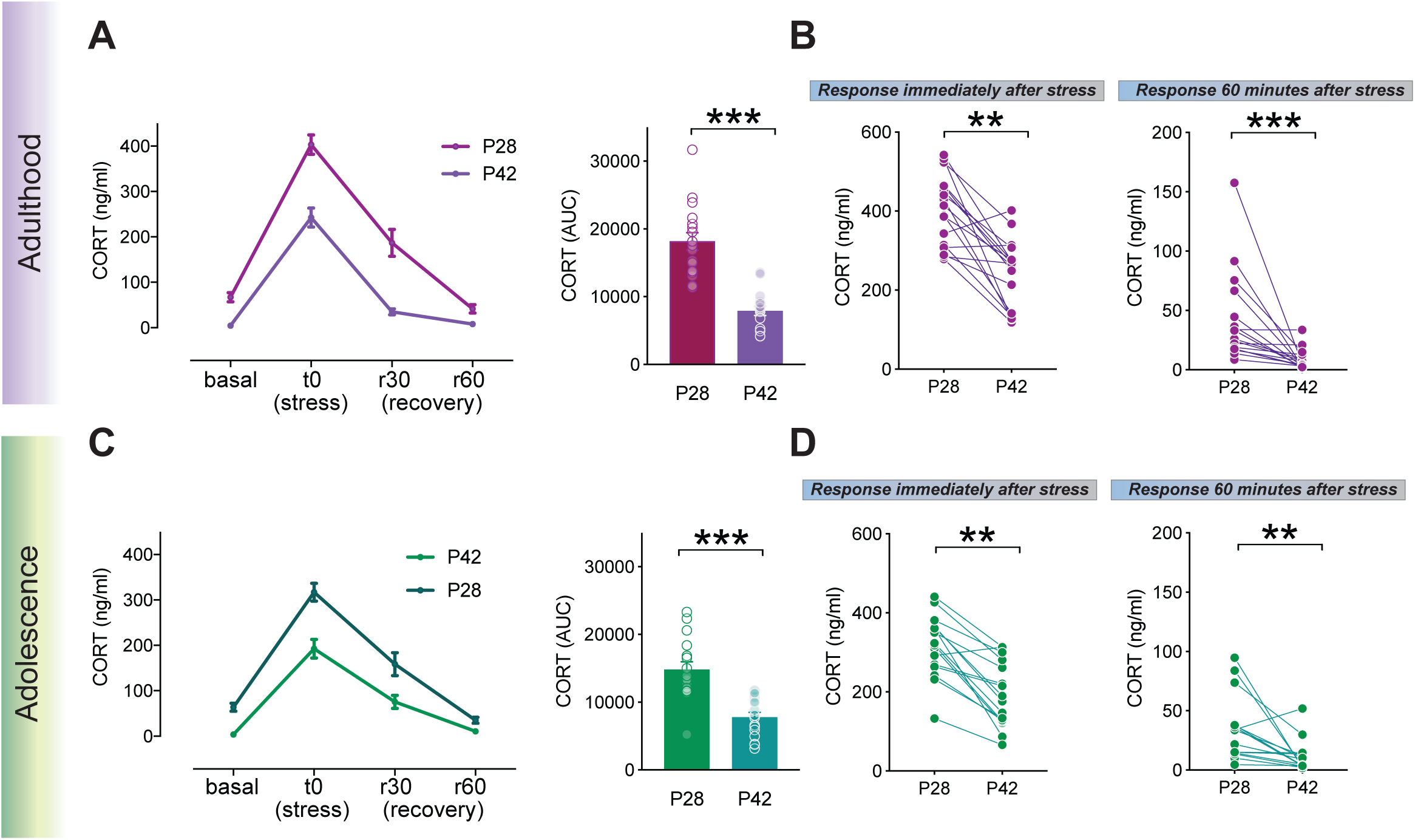
Corticosterone (CORT) response to peripubertal stressor in the two independent batches of animals, where long term effect of PPS was assessed either during Adulthood or Adolescence. The analysis was performed post hoc to examine whether the pattern of corticosterone reactivity to stressors would be the same between the two independent groups. Blood samples were withdrawn during PPS, on P28 and P42 from STRESS rats. **A**. Data from rats assigned to the adulthood group. Left: samples were taken at baseline conditions (i.e. before stressor exposure-basal), immediately following the stressor (t0, stress), as well as, 30 minutes and 60 minutes after the end of the exposure to stress (r30 and r60 respectively). Reduced CORT response was observed on P42 as compared to P28 as confirmed by differences in the AUC (Right). **B**. Individual values for each animal of this batch indicating the CORT response immediately after stress (t0) in P28 and P42 (left) or in the recovery period (r60) (right). **C**. A very similar pattern of corticosterone response was observed in the adolescence group either for the temporal dynamics (left) as for the reduction in the AUC CORT response (right). **D**. Again, marked individual differences in the adaptation to the peak (left) or recovery (right) responses after peripubertal stresss were also observed in the adolescence batch of animals. Results are expressed as the mean ± S.E.M. **p<0.01, ***p<0.001.

